# Oxidative DNA damage is epigenetic by regulating gene transcription via base excision repair

**DOI:** 10.1101/069955

**Authors:** Aaron M. Fleming, Yun Ding, Cynthia J. Burrows

## Abstract

Reactive oxygen species (ROS) have emerged as important cellular signaling agents for survival. Herein, we demonstrate that ROS-mediated oxidation of DNA to yield 8-oxo-7,8-dihydroguanine (OG) in gene promoters is a signaling agent for gene activation. Enhanced gene expression occurs when OG is formed in guanine-rich, potential G-quadruplex sequences (PQS) in promoter coding strands to initiate base excision repair (BER) by 8-oxoguanine DNA glycosylase (OGG1) yielding an abasic site (AP). The AP enables melting of the duplex to unmask the PQS to adopt a G-quadruplex fold in which apurinic/apyrimidinic endonuclease 1 (APE1) binds, but inefficiently cleaves, the AP for activation of *VEGF* or *NTHL1* genes. This concept allowed identification of 61 human DNA repair genes that might be activated by this mechanism. Identification of the oxidatively-modified DNA base OG as guiding protein activity on the genome and altering cellular phenotype ascribes an epigenetic role to OG.

---

Prevailing wisdom suggests oxidatively-induced DNA damage, such as 8-oxo-7,8-dihydroguanine (OG, Fig. 1A), would be detrimental to cellular processes. For example, when OG is located in recognition elements of NF-κB^1^, Sp1^2^, or CREB^3^ transcription factors, protein binding affinity was significantly reduced. When OG was present in the template strand of protein coding regions, modest stalling of RNA pol II occurred^4^, while the initiation of DNA repair at OG to yield an abasic site (AP) stopped RNA pol II leading to truncated transcripts^5^. These observations support a hypothesis of OG decreasing gene transcription. In contrast, mouse livers with infection-induced colitis exhibit increased levels of genomic OG in tandem with enhanced expression of many DNA repair, cell cycle, and stress response genes^6^. Another notable example appears when rat pulmonary artery endothelial cells experience hypoxia; a strong positive correlation between OG in promoter regions and elevated expression of >100 genes was observed^7^. One gene in particular is *VEGF*, for which OG was found in the G-rich potential G-quadruplex sequence (PQS)^7^ that is hypothesized to be responsible for transcriptional regulation of the gene^8^.

**Figure 1.**
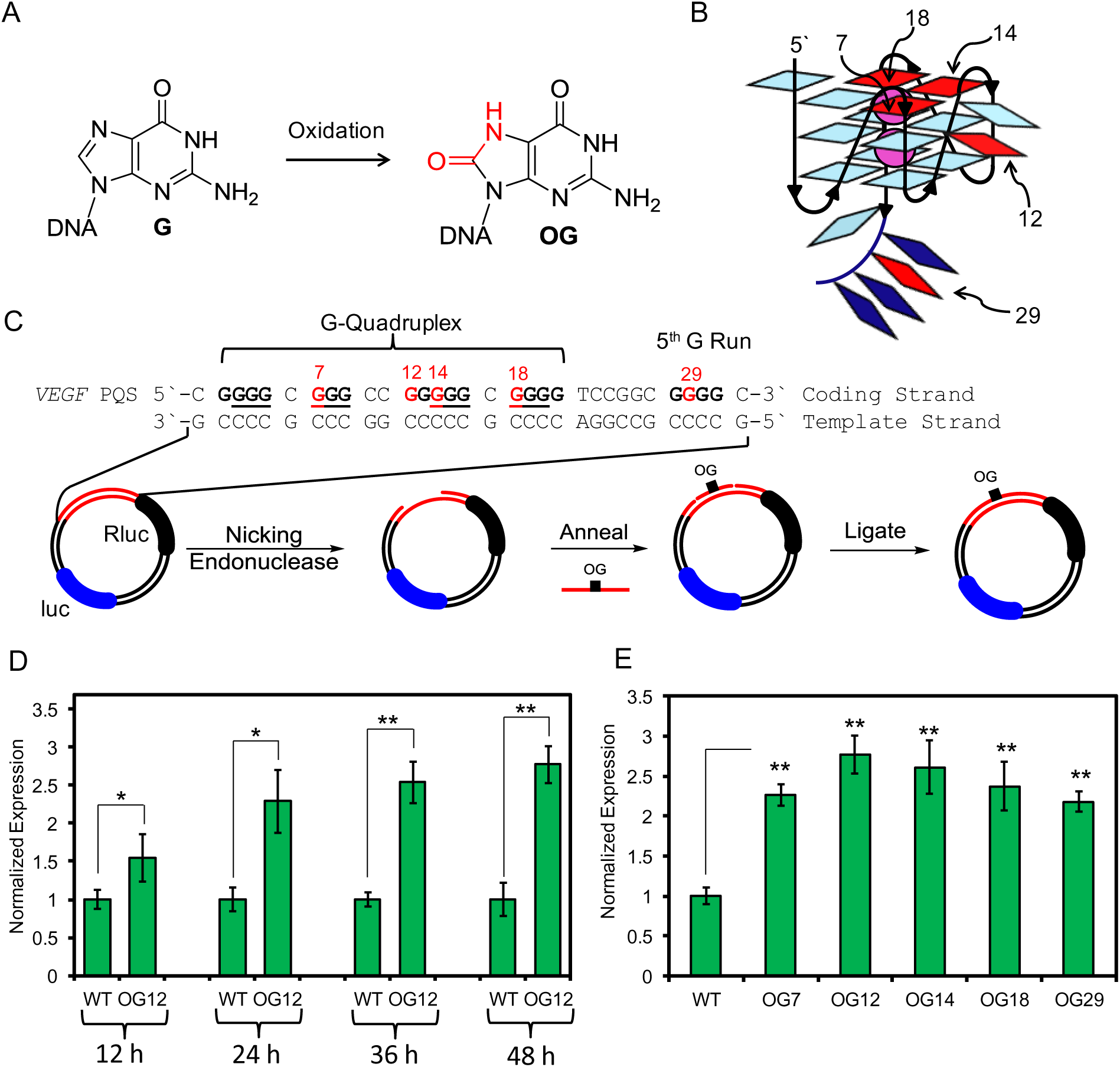
Oxidation of G in the *VEGF* PQS induces transcription. (A) G oxidation to OG. (B) *VEGF* G-quadruplex labeled with positions studied. (C) *VEGF* sequence with Gs in the core underlined, reporter system design, and method for site-specific incorporation of DNA modifications. (D) Time-dependent and (E) position-dependent expression at 48 h post-transfection of OG-containing reporters in glioblastoma cells. Error bars represent 95% c.i., on the basis of 4 or 8 replicates. Significance values for each comparison were calculated by a Student’s t-test. Significance at * *P* < 0.05 or ** *P* < 0.01 is indicated.

The *VEGF* PQS possesses five G tracks, in which the four tracks required for folding to a parallel-stranded G-quadruplex are the 5`-most (Fig. 1B)^9^. Our previous work mapped the most reactive guanine (G) bases in *VEGF* toward oxidation leading to OG and other secondary oxidation products^10^. When DNA damage resides in the four G tracks of the *VEGF* G-quadruplex, DNA repair was not observed^11^; to our surprise, addition of the fifth G track allowed a structural transition to a competent G-quadruplex fold by extruding the damaged G run into a long loop allowing faithful DNA repair^10^. In the present work, an alternative mechanism for gene induction driven by G oxidation to OG that induces a structural switch in the *VEGF* PQS promoter element is proposed and experimentally validated in cells. Induction of transcription was found to require 8-oxoguanine DNA glycosylase (OGG1) and apurinic/apyrimidinic endodeoxyribonuclease 1 (APE1), also known as redox effector factor 1 (Ref-1), in the base excision repair (BER) pathway. Lastly, human DNA repair genes with PQSs also capable of this mechanism were identified, and one was studied, pointing to this mechanism as a possible general approach for up-regulating genes under oxidative stress.

## Results and discussion

Demonstration that OG drives the *VEGF* PQS promoter element to induce transcription was accomplished using a luciferase reporter plasmid. Key features of the reporter system include the *VEGF* PQS promoter element with all five G runs regulating the Renilla luciferase gene (Rluc). The regulatory sequence also includes flanking nicking endonuclease sequences allowing replacement of the G-rich sequence with a synthetic oligomer containing a single, site-specific OG. Additionally, the plasmid possessed the firefly luciferase gene (luc) regulated by an unmodified promoter as an internal standard (Fig. 1C and Fig. S1). The OG positions selected are based on the *VEGF* G-quadruplex structure solved by NMR (Fig. 1B and 1C)^9^.

Changes in gene expression as a function of OG position were studied focusing on the oxidation-prone *VEGF* PQS sites^10^ 7, 12, 14, and 18. Position 12 is in a loop, while 7, 14, and 18 occupy core positions in the G-quadruplex providing contrasting views on DNA damage and structure (Fig. 1B). Position 29 residing in the 5^th^ G track that is not part of the principal G-quadruplex structure was also studied. First, the time-dependent expression of Rluc with OG incorporated at position 12 was evaluated upon transfection into glioblastoma cells, and the reported expression was normalized against the luc expression utilized as an internal standard. From 12 to 48 h post transfection, the expression of Rluc significantly increased to nearly 3-fold when OG was present compared to the WT plasmid (i.e., the all-G-containing plasmid, Fig. 1D). Next, when OG was studied at other sites (7, 14, 18, or 29), measurements made 48 h post transfection enhanced Rluc expression by 2.2- to 3.0-fold (Fig. 1E). These results demonstrate the presence of OG in the *VEGF* promoter PQS enhances the transcriptional output of the reporter gene; significantly, OG was not detrimental, but rather enhanced transcription. This observation is consistent with a previous study monitoring *VEGF* expression under hypoxic conditions that identified a ~3-fold increase in expression^7^.

Experiments to reveal molecular details by which OG induced gene expression were conducted. In the mammalian genome, OG is bound and cleaved by OGG1 in the first step of BER (Fig. 2A). Whether OGG1 cleaves the phosphodiester backbone or APE1 catalyzes this step remains unanswered (Fig. 2A)^12^. To establish if OGG1 is involved in gene induction, comparative studies were conducted in wild-type and OGG1^-/-^ mouse embryonic fibroblasts (MEFs) transfected with OG-containing plasmids. In WT MEFs, depending on the position of OG, Rluc expression increased by 2.5- to 3.9-fold, while OG-containing plasmids in the OGG1^-/-^ MEFs yielded essentially no change in the amounts of Rluc expression compared to the WT plasmid (Fig. 2B). Because OG is in the coding strand of the promoter, it does not interfere with advancement of RNA pol II on the promoter to the transcription start site in the OGG1^-/-^ MEFs, and therefore, the same gene output was observed with and without OG.

**Figure 2.**
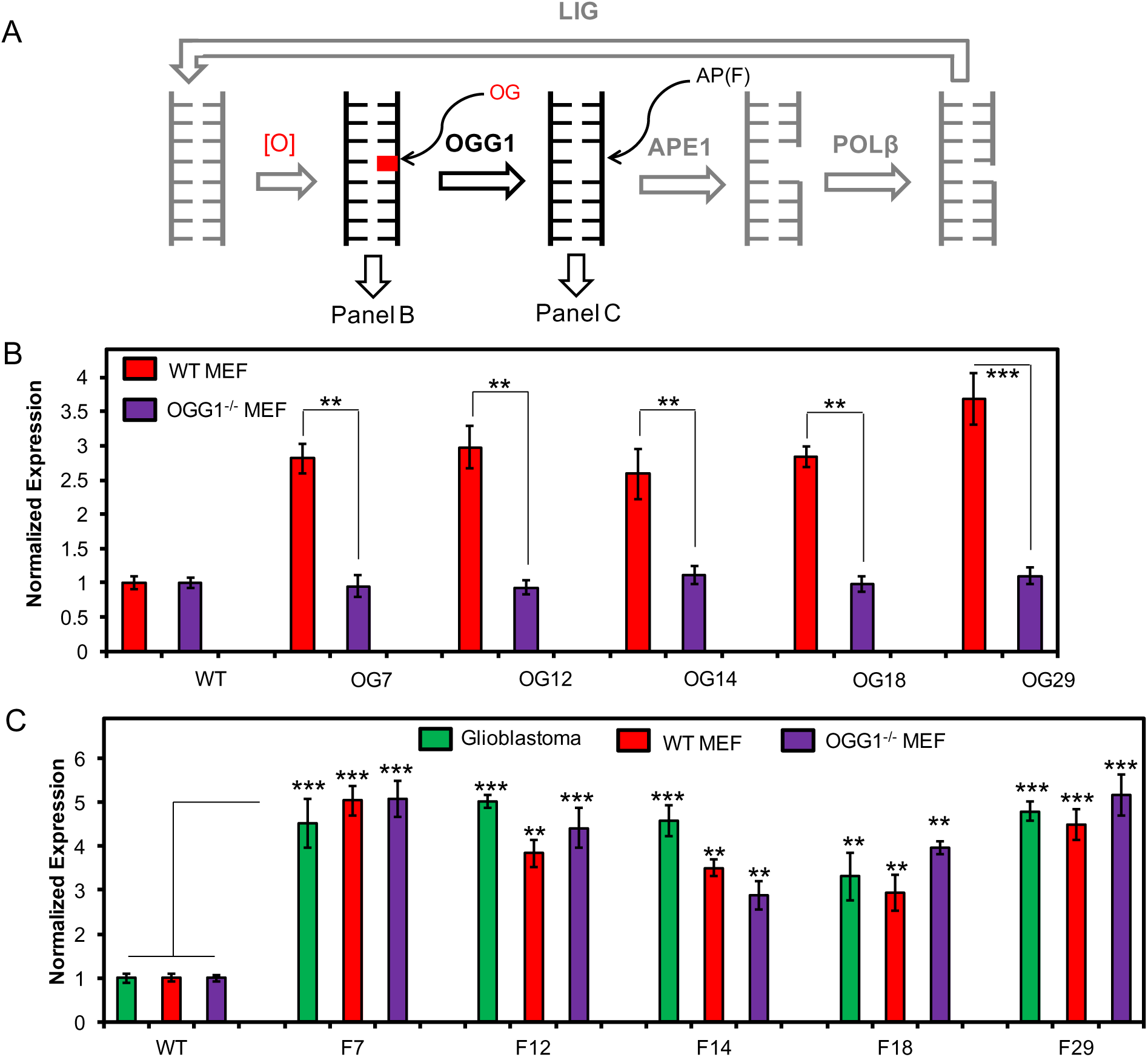
BER is responsible for gene activation when OG is located in a promoter PQS. (A) BER scheme. (B) Positional dependency in expression of OG-containing reporters in WT and OGG1^-/-^ MEFs. (C) Positional and cell line dependency in expression of F-containing reporters, where F = the stable AP analog THF. Error bars represent 95% c.i., on the basis of 4 or 8 replicates. Significance values for each comparison were calculated by a Student’s t-test. Significance at ** *P* < 0.01 or *** *P* < 0.001 is indicated.

The observation of OG not enhancing transcription in the OGG^-/-^ MEFs confirms OGG1 is critical to expression enhancement; however, more questions remain. In wild-type cells, after removal of OG by OGG1, an AP site is present in DNA. Consequently, plasmids with APs (THF analog, F) at the reactive sites were introduced to establish whether gene induction still occurs when the product of OG excision from the DNA is present. Transfection of the AP-containing plasmids into all three cell lines yielded Rluc expression enhancement by 2.5- to 6-fold relative to the WT plasmid (Fig. 2C). These results identify a stronger sequence and cell line dependency in transcriptional amplification than observed with OG. The enhanced expression observed, especially in the OGG1^-/-^ MEFs, identifies APE1 as the possible BER enzyme responsible for increasing gene expression.

To further confirm that APE1 induces transcription when operating on an AP, glioblastoma cells were treated with siRNA specific for knocking down APE1 to prevent binding and cleavage of the AP (Fig. 3A). Glioblastoma cells treated with siRNA were then transfected with plasmids containing an F in the promoter at positions 7, 12, and 29 of the Rluc gene.
Expression of Rluc in the knocked down cell line was similar to the expression of the WT plasmid (Fig. 3B). This initial observation points to the importance of APE1 in gene activation when an AP site is present. Next, we wanted to understand if the ability of APE1 to bind the abasic site or cleave the site was responsible for gene activation (Fig. 3C). First, 1 μM APE1 inhibitor III (Fig. 3D) was added prior to transfection of the F-containing plasmids. The inhibitor prevents cleavage of the backbone while APE1 binding is not strongly impacted (Fig. 3C)^13^. In these studies, Rluc expression increased by >30-fold when an AP was present compared to the WT plasmid (Fig. 3E). For further verification that APE1 binding to APs without causing stand breaks still leads to gene induction, non-cleavable APs were studied (Fig. 3B). Previous experiments found APE1 binds but poorly cleaves oligomers containing a phosphorothioate and 2`-OMe nucleotide 5` to the AP analog F (2`-MeO-PS-F, Fig. 3D)^14^. Therefore, 2`-MeO-PS-F modified plasmids were transfected into glioblastoma cells inducing >10-fold Rluc expression (Fig. 3C). The combined results of these studies support OG inducing transcription via OGG1 generation of an AP in the *VEGF* PQS followed by APE1 binding. More importantly, gene induction occurs while APE1 is bound (Fig. 3C and E), and it does not require the lyase activity of APE1 to yield a strand break for gene activation.

**Figure 3.**
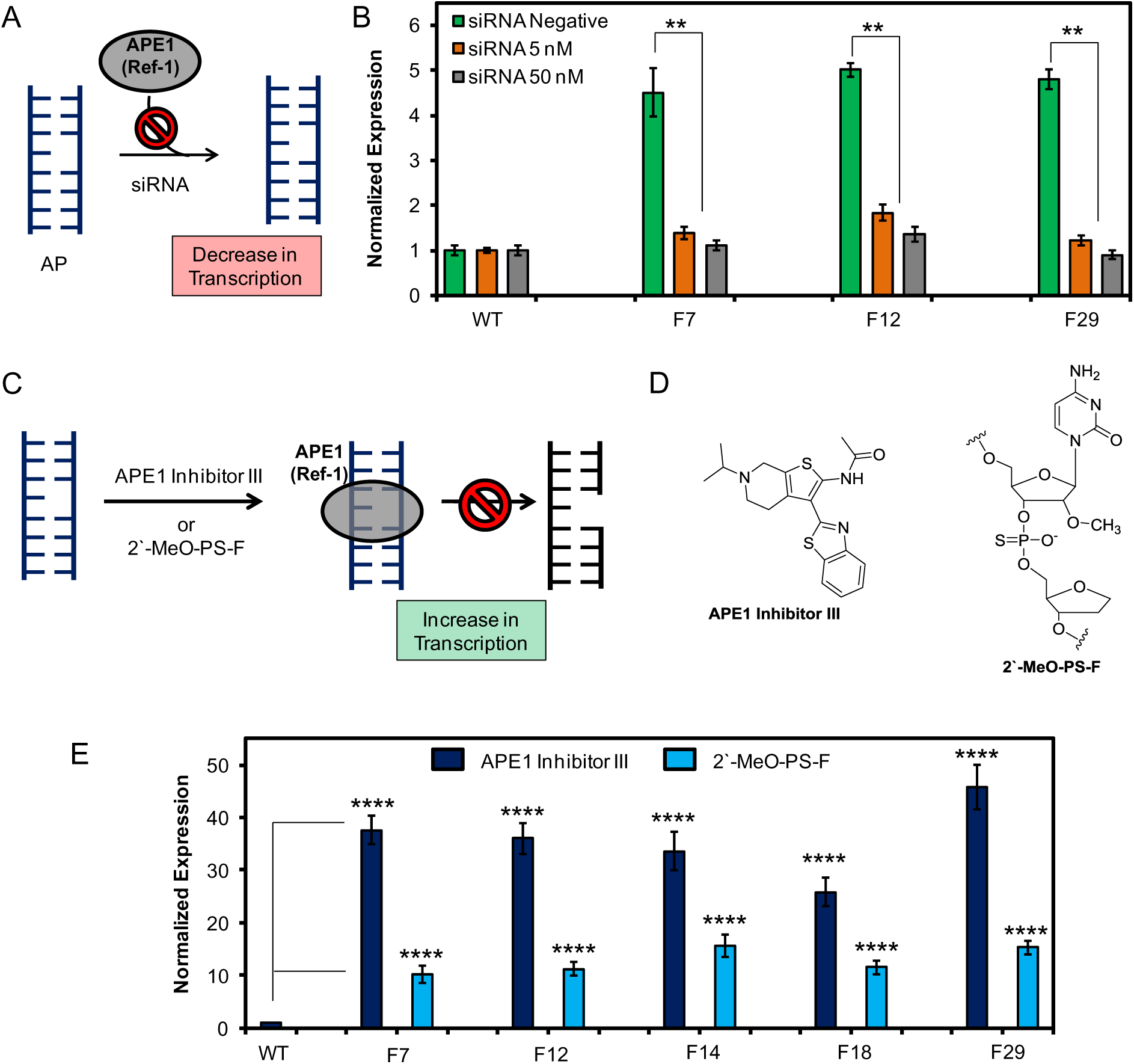
Binding of APE1 (Ref-1) to AP in the *VEGF* PQS promoter element enhances gene transcription in glioblastoma cells. (A) Knock down of APE1 and prevention of AP binding leads to decreased Rluc expression. (B) Expression levels measured when F-containing reporter plasmids were transfected into APE1 siRNA knockdown cells. (C) Mechanism for prevention of APE1 cleavage without impacting binding of an AP leading to increased gene expression. (D) Structures of APE1 inhibitor III and the 2`-MeO-PS-F modified AP site. (E) Expression levels measured when cells were treated with APE1 inhibitor III or they were transfected with poorly cleavable 2`-MeO-PS-F AP. Error bars represent 95% c.i., on the basis of 4 or 8 replicates. Significance values for each comparison were calculated by a Student’s t-test. Significance at ** *P* < 0.01, *** *P* < 0.001, or **** *P* < 0.0001 is indicated.

APE1, also known as redox-effector factor-1 (Ref-1), interacts with many activating transcription factors under oxidative stress conditions leading to enhanced transcription^15^. The link to APE1 (i.e., Ref-1) activation is via OGG1 initiating BER removal of OG to yield an AP site that is bound by APE1. APE1 activates transcription by its Ref-1 activity to guide binding of activating transcription factors. The identity of these activating factors has not yet been pursued. The ability of the VEGF PQS to adopt an alternative fold mediated by the AP site was found to be more critical for activation.

The critical *VEGF* promoter sequence is bound by three Sp1 transcription factors and is a PQS, both of which have been identified as elements in gene activation^8,16^. Delineation of how these two features play into gene induction in the face of DNA damage was pursued. Using a sequence bearing OG yet still recognizable by Sp1 but incapable of G-quadruplex formation, and it was transfected into glioblastoma cells (Fig. S2). Expression of Rluc with the G-quadruplex-negative plasmid containing OG remained similar to expression of the WT plasmid (Fig. 4A). This study confirms that altering Sp1 binding by DNA damage is not critical for induction of transcription; moreover, the G-quadruplex-negative experiment causing no gene induction provides strong support for the G-quadruplex fold as key to gene activation. Further support for considering the G-quadruplex fold in the gene activation mechanism hails from the significant progress demonstrating these folds are critical gene regulatory motifs^17,18^.

**Figure 4.**
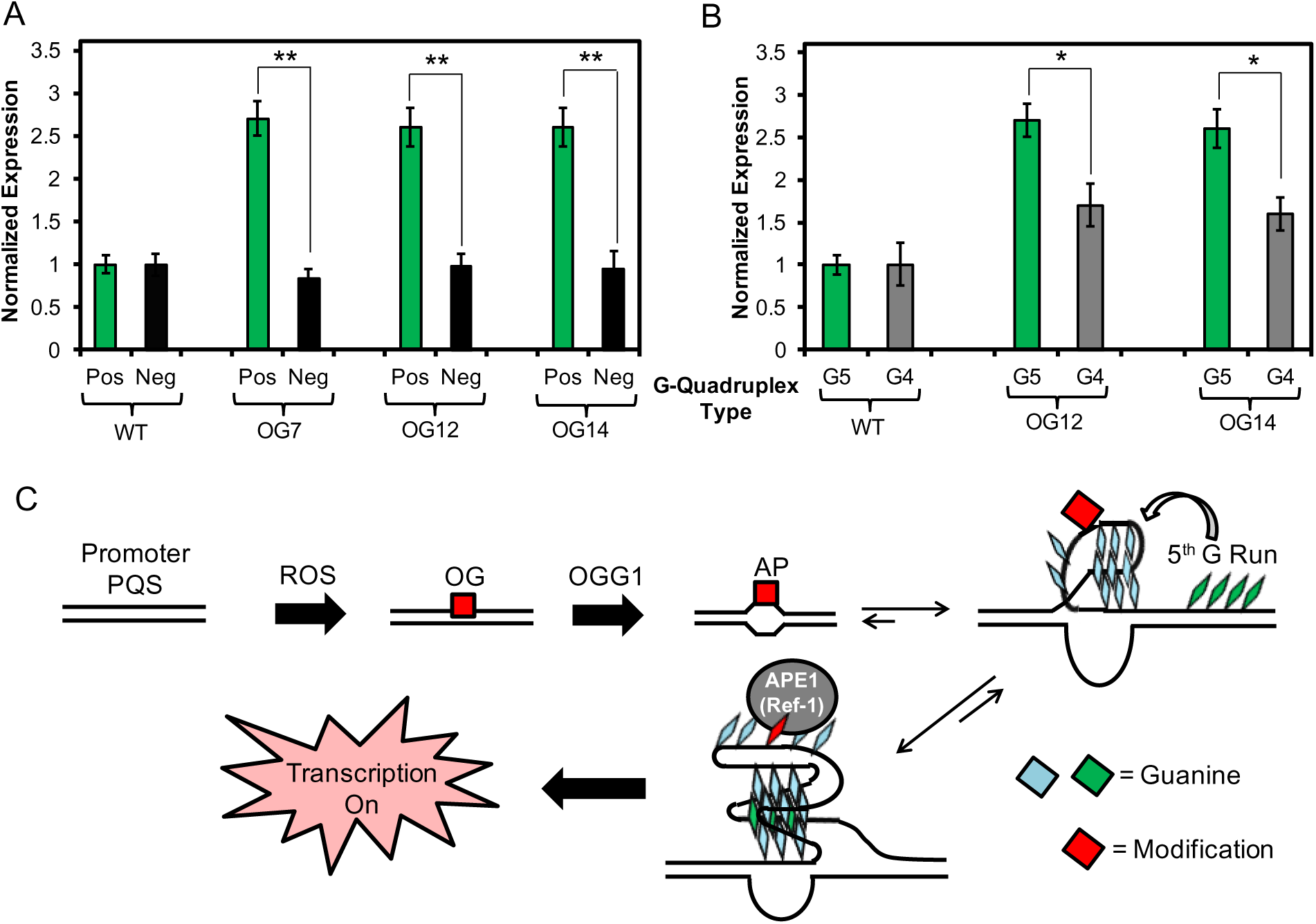
Gene induction with OG in the VEGF PQS requires the G-quadruplex fold. (A) Expression observed when OG is in a G-quadruplex positive or negative folding sequence context (Fig. S2). (B) Requirement of the 5th G run for maximal expression when OG is present. (C) Proposed mechanism for gene induction when OG is present in a promoter PQS. Error bars represent 95% c.i., on the basis of 4 or 8 replicates. Significance values for each comparison were calculated by a Student’s t-test. Significance at * *P* < 0.05 or ** *P* < 0.01 is indicated.

We previously demonstrated the concept that a five track G-quadruplex can switch structures to extrude damaged DNA bases to maintain the alternative fold^10^. To further verify the essential role for the 5^th^ G-track (G5) in gene activation, plasmids containing OG with only the four G runs (G4) required for folding^9^ were transfected. The G4 samples gave significantly less gene expression than the G5 analogs (Fig. 4B); this observation supports a structure-switching gene activation mechanism induced by the 5^th^ G track when DNA damage is present. Thus, gene activation occurs when the sequence adopts a G-quadruplex with the AP extruded into a long loop that is bound by APE1.

On the basis of the proteins involved and the implied requirement of G-quadruplex folding to induce *VEGF* expression when DNA damage is present, the following mechanism is proposed. In the genomic context, the *VEGF* PQS thermodynamically favors the duplex state, in which oxidation of G can be written into the genome directly by (1) cellular oxidants,^6^ (2) via long-range electron transfer through the DNA π stack^19^, or (3) as a result of stress-induced chromatin remodeling (Fig. 4C)^20^. All three oxidation mechanisms will induce G oxidation to OG (Figs. 1A and 4C). In duplex DNA, OG is well accommodated and does not impact duplex stability, although it is readily found and cleaved by OGG1 leading to an AP. The AP decreases the duplex stability by >18°C providing local duplex melting (Fig. 4C and Fig. S3); in contrast, the AP has minimal impact on G-quadruplex stability, because formation of a stable fold can occur via the 5^th^ G-run allowing extrusion of the damaged track into a long loop (Fig. 4C and Fig. S4). Lastly, the AP is strongly bound by APE1 in a G-quadruplex fold, but the cleavage kinetics are attenuated, as previously described^21^. Prolonged APE1 binding induces transcription most likely with the aid of other activating factors.

The sequence of events described provides an alternative mechanism for gene activation under oxidative stress. Is this mechanism specific to *VEGF* or can it be generalized to other genes? Because genomic OG concentrations are elevated in tandem with DNA repair, cell cycle, and stress response genes during inflammation^6^, we inspected 191 human repair and cell cycle genes^22^ against ChIP-Seq data identifying binding sites for the G-quadruplex-specific helicases XPB and XPD^23^. Our analysis of the ChIP-Seq data inspected for promoter PQSs in the DNA repair genes found 61 genes (Fig. 5A and Table S1). We then selected the PQS found in the coding strand of the *NTHL1* promoter for our next study because this gene is also induced under oxidative stress ^24^ and has a 5^th^ G-run allowing structure switching to occur. First, we determined the *NTHL1* PQS could adopt a G-quadruplex and identified sites sensitive to oxidation (Fig. 5B and Fig. S5). Next, the *NTHL1* promoter sequence was introduced into the plasmid system to regulate Rluc expression with OG or F replacing a loop or core reactive G. Transfection of the *NTHL1* plasmid into glioblastoma cells found Rluc expression increased by 4- to 7-fold when the modification was present (Fig. 5C). This further observation confirms these base modifications induce transcription when found in another PQS, supporting this mechanism as a possible general pathway for transcription induction under oxidative stress. The other 60 genes identified are potential subjects of future work, and we note that this list (Fig. 5A) overlaps with those identified by Mangerich et al. as being up-regulated during inflammation and oxidative stress^6^.

**Figure 5.**
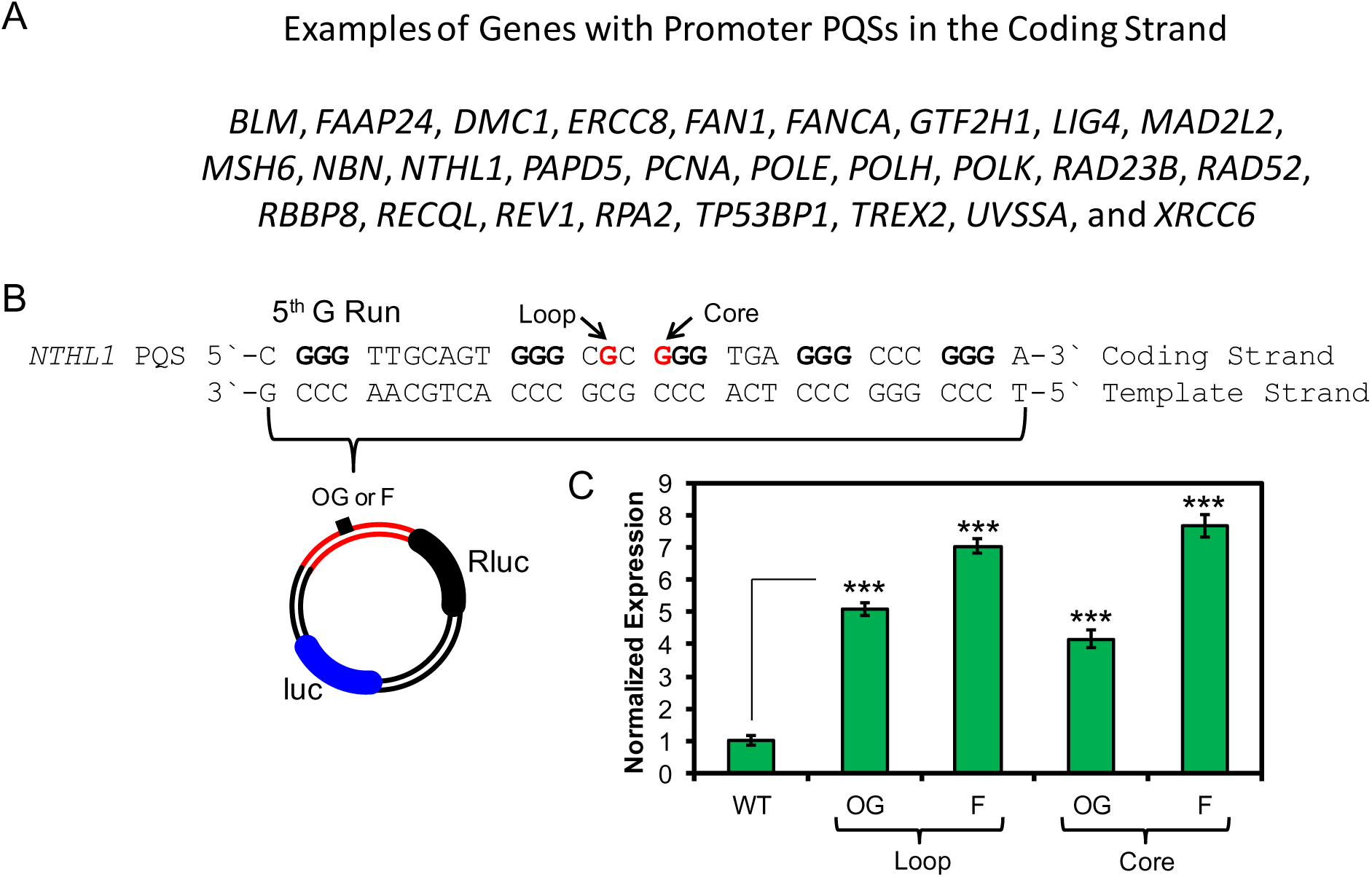
Gene activation is observed when OG or F is present in the *NTHL1* PQS. (A) Partial list of DNA repair genes with coding strand PQSs. (B) The NTHL1 PQS sequence and locations in which OG or F were synthesized. (C) Expression enhancement observed when OG or F is found in the *NTHL1* PQS. Error bars represent 95% c.i., on the basis of 4 or 8 replicates. Significance values for each comparison were calculated by a Student’s t-test. Significance at \*\*\**P*<0.001 is indicated.

The alternative gene activation pathway identified requires base oxidation of G to OG in the PQS context to guide OGG1 for generation of an AP. The AP unmasks the PQS from the duplex state to reveal the G-quadruplex for prolonged binding by APE1 and associated factors (Fig. 4C). The function of oxidative base modifications in DNA to direct proteins to alter transcription ascribes an epigenetic role to OG. The protein readers and erasers are members of the BER pathway, and therefore, these proteins activate genes in addition to guarding the genome against insults such as oxidative stress. Coupling of DNA repair with transcription provides an efficient mechanism to complete two necessary cellular tasks during oxidative stress. An intertwining of these pathways is starting to emerge in other studies^25^. This study demonstrates cells can harness oxidized modifications of DNA bases for altering phenotype under oxidative stress and identifies a mechanism by which ROS are cellular signaling agents, as previously hypothesized^26^. Lastly, the observation that classically defined forms of DNA damage have an epigenetic role in the cell has surfaced with the recent demonstrations of 5-hydroxymethyluracil^27^ and *N*^6^-methyladenine^28^ also guiding cellular processes in higher eukaryotes. These studies extend the epigenetic landscape in DNA beyond methylation of cytosine and its oxidized derivatives^29-31^.

## Methods

Detailed materials and methods are described in Supplementary Section “Materials and methods”. Characterization of the *VEGF* PQS with AP and structural characterization of the *NTHL1* PQS can be found in the Supplementary Section.

**Plasmid construction.** The plasmids were constructed from the psiCHECK2 plasmid (Promega) that contains genes for the Renilla luciferase (Rluc) and firefly luciferase (luc) proteins. The luc gene is regulated by the HSV-TK promoter and used as the internal standard, whereas the Rluc gene was originally regulated by the SV40 early enhancer/promoter that was modified to include the PQS of interest. Additionally, the PQSs of interest were flanked by Nt.BspQ1 nicking endonuclease recognition sequences. Insertion of the PQS and nicking endonuclease recognition sequences was achieved using restriction free cloning, followed by transformation to competent *Escherichia coli* and isolation by miniprep kit (Qiagen), as described previously^32^. The site-specific modifications were synthesized into short oligomers with the sequence between the two nicking endonuclease sites, and they were inserted into the plasmid via literature methods^32,33^. Confirmation that the DNA modifications were introduced into the plasmid was achieved using a protocol established in our laboratory, in which the modification was removed by Fpg and APE1 to yield a ligatable gap^34^. After ligation of the gap with T4-DNA ligase, Sanger sequencing provided a characteristic nucleotide loss at the modification site to confirm its presence.

**Cell Culture Studies.** All cells were grown in Dulbecco’s Modified Eagle Medium (DMEM) supplemented with fetal bovine serum (FBS), gentamicin, glutamax, and non-essential amino acids. The WT MEF and OGG1^-/-^ MEF cells were previously developed and reported on in the literature^35^, and the glioblastoma cells were purchased from ATCC. Transfection experiments were conducted in white, 96-well plates using X-tremeGene HP DNA transfection agent (Roch) with 200 ng of plasmid following the manufacturer’s protocol. The Dual-Glo luciferase (Promega) assay was conducted following the manufacturer’s protocol to monitor Rluc and luc expression levels. Each experiment was conducted in 4 or 8 replicates, as recommended by the Dual-Glo luciferase assay. The APE1 inhibitor III studies were conducted by addition of 1 μM APE1 inhibitor from a DMSO stock solution to the cell culture media during transfection. Controls in which only DMSO was added to the cells found no change in expression level of the wild type plasmid insuring the data obtained resulted from the inhibitor and not the DMSO. The siRNA knockdown studies were conducted with FlexiTube siRNAs (Qiagen) following the manufacturer’s protocol.

**Data analysis.** The data were analyzed by converting the luminescence measured into normalized relative response ratios (RRR), which is the luminescence of Rluc divided by the luminescence of luc (i.e., RRR = Rluc/luc). To obtain the normalized expression values reported, each RRR was divided by the RRR for the wild type sequence in that data set. The error bars represent 95% confidence intervals obtained from the data. Statistical analysis was achieved using a two-tailed Student’s t-test.

## Acknowledgments

The complete methods, data, and PQS sequences for human DNA repair genes are located in the Supplementary Materials. The MEF cell lines were provided by Dr. Tomas Lindahl (Imperial Cancer Research Fund, UK). We thank the University of Utah core facilities for synthesizing the OG- and F-containing oligomers, as well as conducting Sanger sequencing of the plasmids. This work was supported by a National Cancer Institute grant (R01 CA090689).

## Author contributions

A.M.F. and C.J.B conceived the project. A.M.F. and Y.D. performed the research and analyzed the data. A.M.F. and C.J.B. wrote the manuscript.

## Additional information

Supplementary information is available in the online version of the paper. Correspondence and requests for materials should be addressed to C.J.B. or A.M.F.

## References

1. Hailer-Morrison, M.K., Kotler, J.M., Martin, B.D. & Sugden, K.D. Oxidized guanine lesions as modulators of gene transcription. Altered p50 binding affinity and repair shielding by 7,8-dihydro-8-oxo-2′-deoxyguanosine lesions in the NF-kappaB promoter element. Biochemistry 42, 9761–70 (2003).

2. Ramon, O. et al. Effects of 8-oxo-7,8-dihydro-2′-deoxyguanosine on the binding of the transcription factor Sp1 to its cognate target DNA sequence (GC box). Free Radical Res. 31, 217–229 (1999).

3. Moore, S.P., Toomire, K.J. & Strauss, P.R. DNA modifications repaired by base excision repair are epigenetic. DNA Repair (Amst) 12, 1152–1158 (2013).

4. Tornaletti, S., Maeda, L.S., Kolodner, R.D. & Hanawalt, P.C. Effect of 8-oxoguanine on transcription elongation by T7 RNA polymerase and mammalian RNA polymerase II. DNA Repair (Amst) 3, 483–494 (2004).

5. Allgayer, J., Kitsera, N., Bartelt, S., Epe, B. & Khobta, A. Widespread transcriptional gene inactivation initiated by a repair intermediate of 8-oxoguanine. Nucleic Acids Res., 10.1093/nar/gkw473 (2016).

6. Mangerich, A. et al. Infection-induced colitis in mice causes dynamic and tissue-specific changes in stress response and DNA damage leading to colon cancer. Proc. Natl. Acad. Sci. U.S.A. 109, E1820–E1829 (2012).

7. Pastukh, V. et al. An oxidative DNA “damage” and repair mechanism localized in the VEGF promoter is important for hypoxia-induced VEGF mRNA expression. Am. J. Physiol. Lung Cell Mol. Physiol. 309, L1367–1375 (2016).

8. Sun, D. et al. The proximal promoter region of the human vascular endothelial growth factor gene has a G-quadruplex structure that can be targeted by G-quadruplex-interactive agents. Mol. Cancer Ther. 7, 880–889 (2008).

9. Agrawal, P., Hatzakis, E., Guo, K., Carver, M. & Yang, D. Solution structure of the major G-quadruplex formed in the human VEGF promoter in K^+^: insights into loop interactions of the parallel G-quadruplexes. Nucleic Acids Res. 41, 10584–10592 (2013).

10. Fleming, A.M., Zhou, J., Wallace, S.S. & Burrows, C.J. A role for the fifth G-track in G-quadruplex forming oncogene promoter sequences during oxidative stress: Do these “spare tires” have an evolved function? ACS Cent. Sci. 1, 226–233 (2015).

11. Zhou, J., Fleming, A.M., Averill, A.M., Burrows, C.J. & Wallace, S.S. The NEIL glycosylases remove oxidized guanine lesions from telomeric and promoter quadruplex DNA structures. Nucleic Acids Res. 43, 4039–4054 (2015).

12. Wallace, S.S. Base excision repair: A critical player in many games. DNA Repair 19, 14-26 (2014).

13. Rai, G. et al. Synthesis, biological evaluation, and structure-activity relationships of a novel class of apurinic/apyrimidinic endonuclease 1 inhibitors. J. Med. Chem. 55, 3101–3112 (2012).

14. Freudenthal, B.D., Beard, W.A., Cuneo, M.J., Dyrkheeva, N.S. & Wilson, S.H. Capturing snapshots of APE1 processing DNA damage. Nat. Struct. Mol. Biol. 22, 924–931 (2015).

15. Thakur, S., Dhiman, M., Tell, G. & Mantha, A.K. A review on protein-protein interaction network of APE1/Ref-1 and its associated biological functions. Cell Biochem. Funct. 33, 101–112 (2015).

16. Schafer, G. et al. Oxidative stress regulates vascular endothelial growth factor-A gene transcription through Sp1- and Sp3-dependent activation of two proximal GC-rich promoter elements. J. Biol. Chem. 278, 8190–8198 (2003).

17. Bochman, M.L., Paeschke, K. & Zakian, V.A. DNA secondary structures: stability and function of G-quadruplex structures. Nat. Rev. Genet. 13, 770–780 (2012).

18. Balasubramanian, S., Hurley, L.H. & Neidle, S. Targeting G-quadruplexes in gene promoters: A novel anticancer strategy? Nat. Rev. Drug Discov. 10, 261–275 (2011).

19. Genereux, J.C. & Barton, J.K. Mechanisms for DNA charge transport. Chem. Rev. 110, 1642–1662 (2010).

20. Perillo, B. et al. DNA oxidation as triggered by H3K9me2 demethylation drives estrogen-induced gene expression. Science 319, 202–206 (2008).

21. Broxson, C., Hayner, J.N., Beckett, J., Bloom, L.B. & Tornaletti, S. Human AP endonuclease inefficiently removes abasic sites within G4 structures compared to duplex DNA. Nucleic Acids Res. 42, 7708–7719 (2014).

22. Wood, R.D., Mitchell, M., Sgouros, J. & Lindahl, T. Human DNA repair genes. Science 291, 1284–1289 (2001).

23. Gray, L.T., Vallur, A.C., Eddy, J. & Maizels, N. G quadruplexes are genomewide targets of transcriptional helicases XPB and XPD. Nat. Chem. Biol. 10, 313–318 (2014).

24. Hironaka, K., Factor, V.M., Calvisi, D.F., Conner, E.A. & Thorgeirsson, S.S. Dysregulation of DNA repair pathways in a transforming growth factor alpha/c-myc transgenic mouse model of accelerated hepatocarcinogenesis. Lab Invest. 83, 643–654 (2003).

25. Fong, Y.W., Cattoglio, C. & Tjian, R. The intertwined roles of transcription and repair proteins. Mol. Cell 52, 291–302 (2013).

26. Trachootham, D., Lu, W., Ogasawara, M.A., Nilsa, R.D. & Huang, P. Redox regulation of cell survival. Antioxid. Redox Signal 10, 1343–1374 (2008).

27. Pfaffeneder, T. et al. Tet oxidizes thymine to 5-hydroxymethyluracil in mouse embryonic stem cell DNA. Nat. Chem. Biol. 10, 574–581 (2014).

28. Zhang, G. et al. N(6)-methyladenine DNA modification in Drosophila. Cell 161, 893–906 (2015).

29. Chen, K., Zhao, Boxuan S. & He, C. Nucleic acid modifications in regulation of gene expression. Cell Chem. Biol. 23, 74–85 (2016).

30. Booth, M.J., Raiber, E.-A. & Balasubramanian, S. Chemical methods for decoding cytosine modifications in DNA. Chem. Rev. 115, 2240–2254 (2015).

31. Wagner, M. et al. Age-dependent levels of 5-methyl-, 5-hydroxymethyl-, and 5-formylcytosine in human and mouse brain tissues. Angew. Chem. Int. Ed. Engl. 54, 12511–12514 (2015).

32. Riedl, J., Ding, Y., Fleming, A.M. & Burrows, C.J. Identification of DNA lesions using a third base pair for amplification and nanopore sequencing. Nat. Commun. 6, 8807 (2015).

33. You, C. et al. A quantitative assay for assessing the effects of DNA lesions on transcription. Nat. Chem. Biol. 8, 817–822 (2012).

34. Riedl, J., Fleming, A.M. & Burrows, C.J. Sequencing of DNA lesions facilitated by site-specific excision via base excision repair DNA glycosylases yielding ligatable gaps. J. Am. Chem. Soc. 138, 491–494 (2015).

35. Klungland, A. et al. Accumulation of premutagenic DNA lesions in mice defective in removal of oxidative base damage. Proc. Natl. Acad. Sci. U. S. A. 96, 13300–13305 (1999).

